# Ancient Rapid Radiation Underlies Persistent Phylogenomic Conflict in Early Collembola Diversification

**DOI:** 10.64898/2026.07.05.736609

**Authors:** Claudio Cucini, Edmund R R Moody, Stephen H Montgomery, Francesco Cicconardi

## Abstract

Collembola (springtails) are among the most abundant and ecologically important soil arthropods, representing one of the oldest extant terrestrial hexapod lineages, with a fossil record extending to the early Devonian. Despite their relevance, phylogenetic relationships among the four extant orders (Entomobryomorpha, Poduromorpha, Symphypleona, and Neelipleona) have remained unresolved for over two decades. Here, we present the most comprehensive phylogenomic analysis of Collembola to date, comprising 1,127 single-copy orthologues from 145 taxa representing 19 families. To improve orthology inference, we developed a novel HMM-based filtering pipeline that significantly reduced hidden paralogy in BUSCO-derived datasets. Across multiple dataset configurations, gene-jackknife replicates, and various maximum-likelihood analyses, we consistently recovered Poduromorpha as the earliest-diverging lineage. Coalescent-based methods instead highlighted discordant arrangements characterised by extremely short internal branches and low quartet support, a pattern consistent with pervasive incomplete lineage sorting and reticulate evolutionary history. We further dissected the phylogenetic signal by exhaustively evaluating all possible inter-order topological arrangements, both on the full concatenated dataset and gene-by-gene, to identify the most phylogenetically informative loci. These analyses rejected the great majority of previously proposed hypotheses, narrowing support to only two statistically indistinguishable topologies (T11 and T4), with the Poduromorpha-first arrangement consistently favoured across both site-homogeneous and site-heterogeneous substitution models. Finally, with molecular dating, we estimated the origin of crown Collembola in the Early Devonian, with the diversification of the extant orders in the Carboniferous. Several extant genera were estimated to be older than many currently recognized families, highlighting the exceptional evolutionary persistence of springtail lineages and suggesting that lineage longevity should be considered when interpreting higher-level taxonomic diversity.

---

Collembola, commonly known as the springtails, are amongst the most abundant and ecologically significant arthropods in terrestrial ecosystems. They play key roles in organic matter decomposition, regulation of fungal communities, soil structure formation, and nutrient cycling across virtually all terrestrial biomes (Potapov, et al. 2020, Cucini, et al. 2021), making them valuable bioindicators (Potapov, et al. 2023). Despite there being nearly 9,000 formally described species, molecular studies have repeatedly revealed extensive cryptic diversity (Cicconardi, et al. 2010, Emerson, et al. 2011, Carapelli 2020), and global estimates suggest that the true number of species may exceed 50,000 (Cicconardi, et al. 2013). Collembola therefore remain one of the largest animal radiations without a stable phylogenetic framework.

Together with Protura and Diplura, Collembola belong to the Entognatha, and constitute one of the earliest branching hexapod lineages (Tihelka, et al. 2021), although their phylogenetic position relative to insects remains a subject of active investigation and debate (Du, et al. 2024, Machida, et al. 2025). Their fossil record extends back to the Early Devonian, with *Rhyniella praecursor* from the Rhynie Chert of Scotland (~400 Mya) representing the oldest unambiguous collembolan and, consequently, one of the oldest known hexapods (Whalley and Jarzembowski 1981, Greenslade and Whalley 1986). This ancient evolutionary history makes Collembola a key lineage for understanding the early diversification of terrestrial hexapods and the evolution of soil ecosystems.

The classification of Collembola has undergone substantial revision over the last century. Early classification divided springtails into the elongate-bodied Arthropleona and the globular-bodied Symphypleona *sensu lato* (Borner 1901), a framework that dominated collembolan systematics throughout much of the twentieth century (Fig. 1a). Subsequent morphological and molecular studies suggested that Arthropleona is not monophyletic, leading to the currently recognized subdivision into four orders: Entomobryomorpha, Poduromorpha, Symphypleona *sensu stricto*, and Neelipleona (Deharveng 2004). Despite broad acceptance of this classification, the relationships amongst these four lineages remains a persistent unresolved problem in springtail systematics and one of the most challenging questions in deep hexapod phylogenetics.

**Figure 1.**
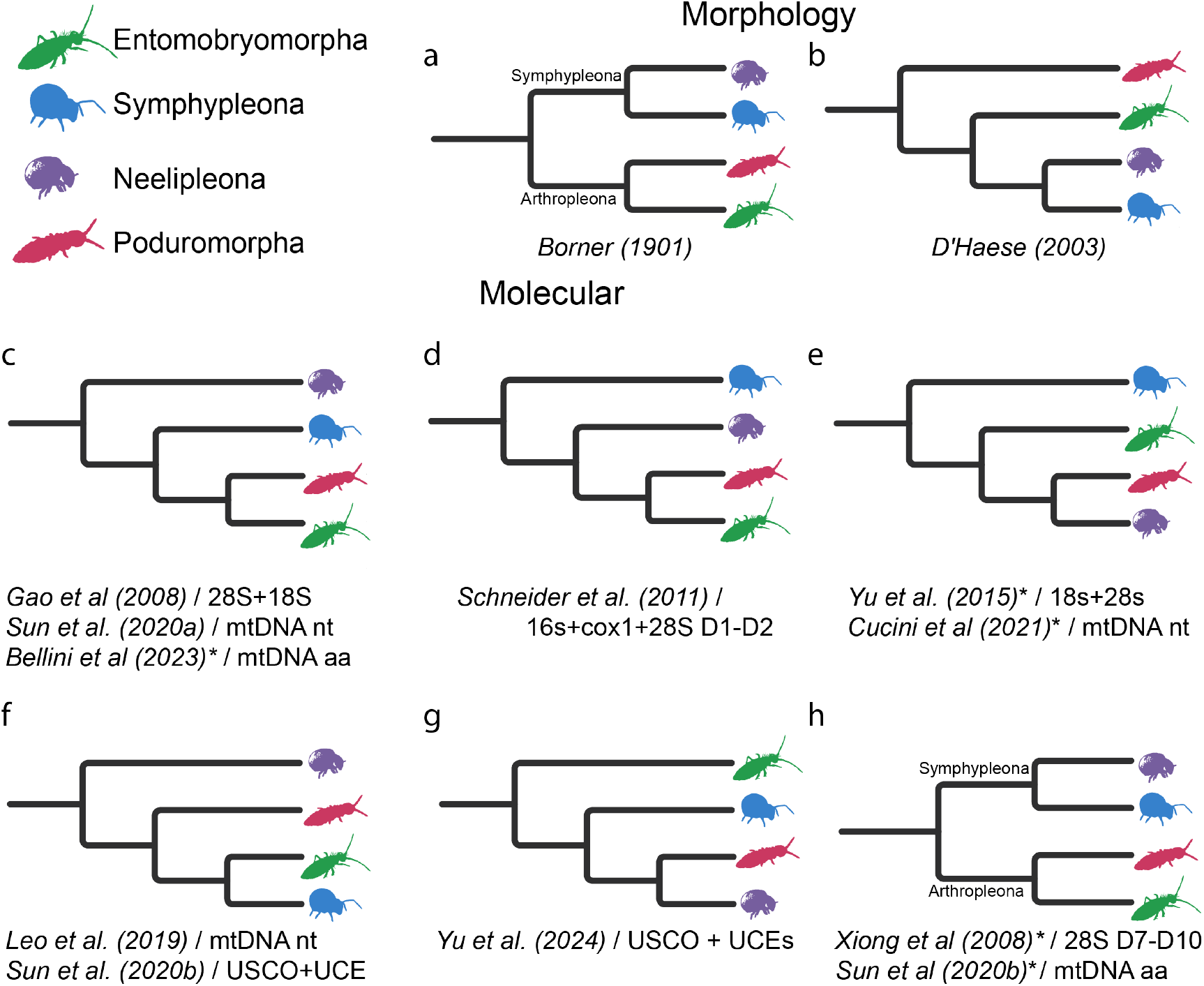
Historical hypotheses of the relationships among the four extant orders of Collembola, based on morphological (**a-b**) and molecular (**c-g**) studies that sampled at least one representative of each of the four orders. Asterisks (*) denote topologies simplified from the original publication.

Over the past two decades, successive generations of phylogenetic datasets have produced conflicting hypotheses regarding the inter-order relationships. Morphological analyses (D’Haese 2003), ribosomal markers (Gao, et al. 2008, Yu, et al. 2016), multilocus datasets (Xiong, et al. 2008, Schneider, et al. 2011), mitogenomes (Leo, et al. 2019, Sun, et al. 2020a, Cucini, et al. 2021, Potapov, et al. 2023), and more recently phylogenomic datasets (Sun, et al. 2020b, Yu, et al. 2024) have each supported alternative branching arrangements of the four clades, often depending on the employed analytical framework; highlighting the difficulty of resolving their phylogenetic relations (Fig. 1).

Although phylogenomics studies have improved resolution, genomic sampling remains remarkably sparse, with only a small fraction of described collembola diversity represented by genome-scale datasets (Cucini, et al. 2025). Consequently, deep divergence within Collembola remains sensitive to both taxon sampling and analytical methodology. Resolving these relationships would also provide a framework for investigating the evolution of key ecological and genomic innovations during the early diversification of terrestrial hexapods. More broadly, Collembola are an excellent model for understanding why some ancient radiations remain resistant to phylogenomic resolution despite genome-scale data.

Here, we reconstruct the most comprehensive phylogenomic framework for Collembola to date using more than 1,100 curated single-copy orthologues sampled across 145 taxa representing all four orders and 19 families. Combining newly generated and publicly available genomes, we integrate concatenation, coalescent, concordance, topology-testing, and divergence-time analyses to evaluate all competing hypotheses of ordinal relationships and investigate the evolutionary processes underlying persistent phylogenetic conflict. Together, these analyses provide a robust phylogenomic framework for Collembola and shed light on the mechanisms responsible for one of the oldest unresolved radiations among terrestrial hexapods.

## Materials AND Methods

### Single-copy orthologue selection and gene-tree inference

To maximize taxonomic sampling, we combined publicly available genomes with draft assemblies from the MetaInvert resource (Collins, et al. 2023), which substantially expanded representation across soil arthropods using a genome-skimming approach, albeit with more fragmented genomes as a trade-off (accessed January 2023) (Table S1). Single-copy orthologous groups were recovered using a BUSCO-based orthology inference pipeline, providing a standardized and computationally efficient framework for orthology inference across large taxonomic scales (Alam, et al. 2025). The accuracy of BUSCO-based approaches can be undermined by fragmented assemblies which frequently contain incomplete genes, split gene models, or incorrectly assigned paralogues, increasing the likelihood of hidden paralogy and potentially biasing downstream phylogenetic inference. To mitigate these effects, we developed an HMM-based filtering pipeline that uses the BUSCO profile HMMs to identify hidden paralogy and domain-level misassignments, while allowing duplicated loci to be partitioned into independent copies when supported by the data, which were treated as independent orthologous partitions (designated A and B) for all downstream analyses (Fig. 2).

**Figure 2.**
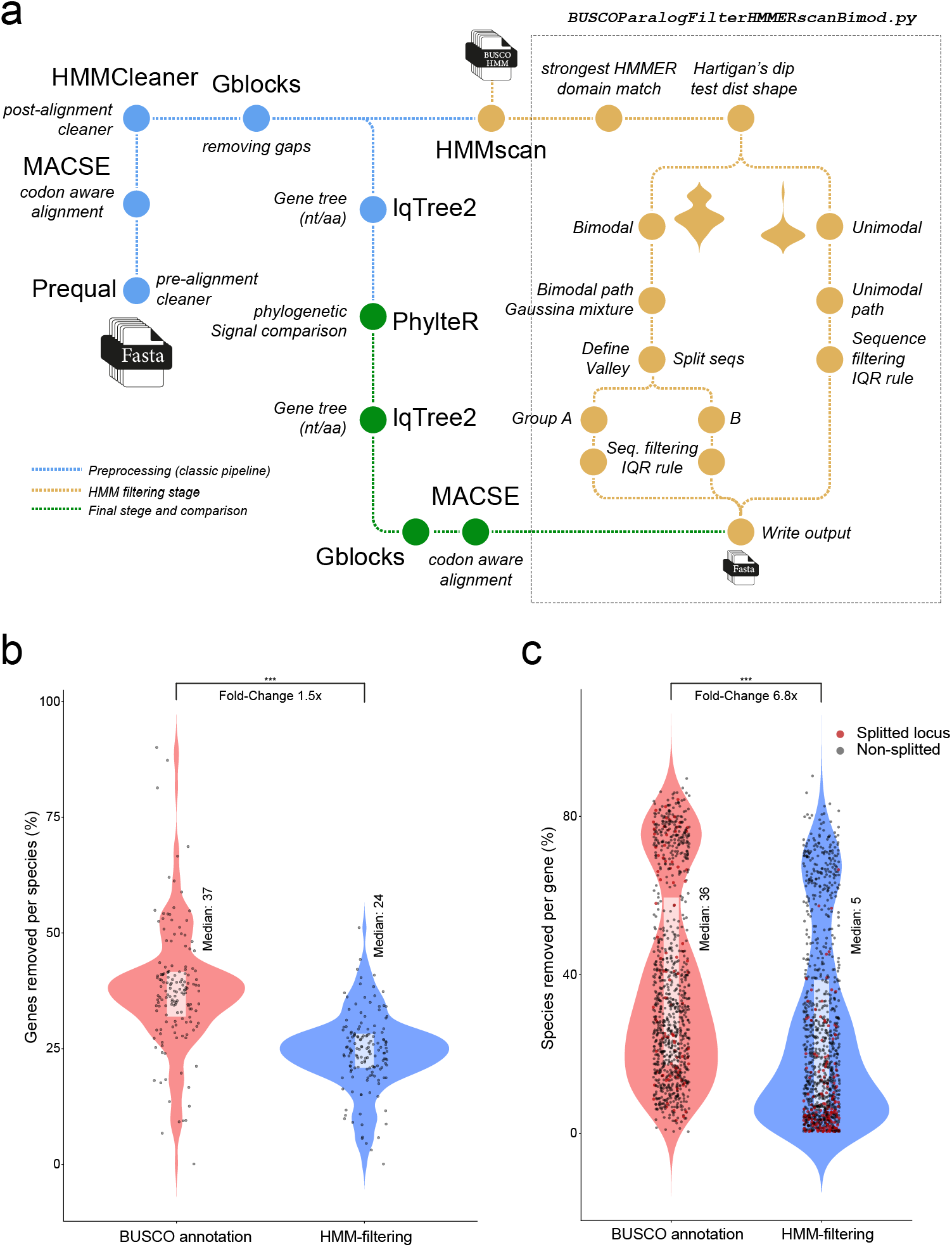
HMM-based filtering reduces hidden paralogy at both taxon and locus levels. **a**) Schematic overview of the BUSCO paralog filtering pipeline. The pipeline outputs filtered FASTA files and corresponding outlier lists, retaining sequences with consistent HMMER support while excluding outliers, sequences that are too divergent or weak matches. **b**) Distributions of the proportion of genes removed per species before (classic quality-filter compleASM BUSCO annotation) and after HMM-based filtering pipeline (yellow in A). In both panels, violin plots show the distribution of values, while boxplots indicate medians and interquartile ranges, and points represent individual loci or species. Red dots indicate loci from a binomial distribution that were split into two independent sets. Statistical significance was assessed using Wilcoxon tests (*** p < 0.001).

For each assembly, single-copy orthologous genes were identified and annotated using Compleasm V0.2.4 (Huang and Li 2023) with BUSCO gene sets from the Arthropoda lineage dataset odb10 (Waterhouse, et al. 2018). Single-copy orthologous groups were subsequently quality-filtered following the established pipeline used in several other studies (Cicconardi, et al. 2017, Cicconardi, et al. 2020, Cicconardi, et al. 2023, Cicconardi, et al. 2024). Briefly, nucleotide coding sequences were first normalised to preserve reading frame integrity and PREQUAL V1.02 (Whelan, et al. 2018) was used to remove erroneous sequences prior to alignment. Codon-aware alignments were then generated with MACSE V2.03 using the BLOSUM62 scoring matrix (Ranwez, et al. 2018), and further cleaned with HmmCleaner (Di Franco, et al. 2019) to remove compositionally or phylogenetically suspicious regions. Cleaned sequences were realigned with MACSE, and poorly conserved positions were trimmed using Gblocks V0.91b (Castresana 2000) with codon-aware settings and relaxed filtering criteria, requiring approximately half of the sequences plus one to define conserved blocks. If Gblocks returned an empty alignment, trimAl 1.4.REV22 (Capella-Gutierrez, et al. 2009) was then adopted using the gappyout option.

At this stage, we implemented a novel filtering strategy in a python script (BUSCOParalogFilterhmmerscanBimod), available at the Zenodo repository. The code uses the hidden Markov model (HMM) profile associated with each BUSCO ortholog to further reduce phylogenetic noise, with particular focus on hidden paralogy and domain-level misidentification. Each BUSCO gene was screened with HMMER v3.3.2 (Eddy 2011) against its own BUSCO HMM profile, and the resulting HMMSCAN domtblout files were parsed with the best conditional E-value (c-Evalue) retained for each sequence in the orthologous group. Values were then log-transformed, with zero E-values replaced by the smallest non-zero E-value observed in that locus before transformation. To identify loci containing more than one divergent sequence class, we tested the distribution of log-transformed best c-Evalues using Hartigan’s dip test. Bimodal score distributions are expected when orthologous groups contain distinct sequence classes, such as true orthologs and divergent paralogs or partial domain matches. When significant bimodality was detected (P < 0.001), a two-component Gaussian mixture model was fitted, and sequences split into two groups using the midpoint between the two component means. For loci showing no evidence of bimodality, all sequences were analysed as a single group. Within each resulting sequence set, outliers were identified using the interquartile-range (IQR) criterion. Sequences with log-transformed best c-Evalues outside the interval Q1 − 1.5 × IQR to Q3 + 1.5 × IQR were excluded. The significance threshold for the dip test and the IQR were selected following inspection of numerous orthologous groups to provide a conservative and consistent filtering framework across all loci. Filtered FASTA files were generated for downstream alignment and phylogenomic analyses, and excluded sequence identifiers were recorded for each locus. Diagnostic violin and strip plots were generated to inspect the score distributions, mixture-model splits, and IQR thresholds.

Gene trees were inferred from both the initial quality-filtered alignments and the realigned post-filtering datasets, using IQ-TREE v2.1.3 covid-edition (Minh, et al. 2020). For nucleotide alignments, ModelFinder was used to select the best-fitting substitution model. For amino-acid alignments, ModelFinder was run with an expanded candidate set including empirical profile-mixture models (C10-C60) and mixture models combining LG or WAG exchangeabilities with C10-C60 profile components (LG+C10-C60+F and WAG+C10-C60+F). The expanded model set was included to account for across-site amino-acid compositional heterogeneity, which has been shown to improve phylogenetic inference in deep evolutionary datasets (Blanquart and Lartillot 2008). Branch support was assessed using 5,000 ultrafast bootstrap replicates and 5,000 SH-aLRT replicates. Identical sequences were retained during tree inference. To assess whether HMM-based filtering reduced hidden paralogy, gene trees inferred before and after BUSCO-HMM filtering were analysed with PhylteR V0.9.12 (Comte, et al. 2023) to evaluate whether the filtering procedure improved phylogenetic signal by reducing artefacts consistent with hidden paralogy or contamination.

### Concatenated-based phylogenomic analyses

Phylogenetic analyses were conducted using both concatenation- and coalescent-based approaches to evaluate the robustness of relationships across Collembola and to assess the effects of missing data, compositional heterogeneity, locus sampling, and analytical framework on phylogenetic inference. Concatenated maximum-likelihood (ML) analyses were performed using IQ-TREE2 on both nucleotide (FullNTsmatrix) and amino-acid (FullAAsmatrix) supermatrices. Amino-acid analyses were conducted under a gene-wise partitioned LG+F+Γ model, whereas nucleotide analyses used ModelFinder (-m MFP) to select the best-fitting substitution model for each partition. Because inference under empirical profile-mixture models (LG+C10-C60+F+Γ) was computationally prohibitive for the complete supermatrices, increasingly complex mixture models were instead applied to reduced datasets and in formal topology tests to evaluate the robustness of the inferred ordinal relationships to model complexity (see below).

The robustness of phylogenetic inference to missing data was evaluated by analysing datasets filtered to retain loci present in at least 50%, 70%, and 75% of taxa, on both amino acid (model: LG+F+Γ) and nucleotide (GTR+F+Γ10) data type. To evaluate the sensitivity to locus sampling, 20 gene-jackknife replicates were generated by randomly sampling 50%, 70%, and 90% of loci from the complete amino-acid dataset (JK50pAA; JK70pAA; JK90pAA). Each replicate was analysed under LG+F+Γ, and the resulting topologies were compared to assess phylogenetic stability. Finally, an additional amino-acid analysis excluding Neelipleona was performed to evaluate the potential influence of long branch attraction (FullAAsmatrix.NoNeelipleona).

To assess the potential impact of compositional heterogeneity, amino-acid alignments were recoded using the Dayhoff-6 and SR4 reduced-state alphabets (Dayhoff, et al. 1978, Susko and Roger 2007) using RECODING_ALIGNMENTS.R available at https://github.com/HS6986/recoded-mixture-models-new. Dayhoff-6 and SR4 were used to evaluate whether substantial reduction in amino-acid state space affected the inferred topology. The use of two independent recoding schemes allowed us to distinguish topology changes associated with reduced compositional heterogeneity from those specific to a particular recoding strategy. Recoded alignments were analysed in IQ-TREE2 on both the complete dataset and on datasets filtered to retain sites with at least 70% and 75% occupancy (70pOccAA; 75pOccAA), using models appropriate to the recoded character type. Dayhoff-6 recoded matrices were treated as morphological multistate data and analysed under the MK+Γ model (-st MORPH -m MK+Γ), whereas SR4 recoded matrices were treated as four-state DNA-like data and analysed under GTR+Γ (-st DNA -m GTR+Γ). Branch support was assessed using 1,000 ultrafast bootstrap replicates and 1,000 SH-aLRT replicates. All datasets and analysis acronyms can be found at table S4.

### Coalescent species-tree inference and concordance analyses

Phylogenetic conflict was investigated using complementary gene tree-, site-, and multispecies coalescent-based approaches. Species trees were inferred under the multispecies coalescent using ASTRAL V5.6.3 (Zhang, et al. 2018). Analyses were performed separately on amino acid and nucleotide gene trees, including both complete datasets and occupancy-filtered datasets (50%, 70%, and 75% occupancy thresholds). Gene-jackknife replicate trees were also analysed with ASTRAL to evaluate the effects of locus sampling on species-tree inference. To quantify phylogenetic conflict and concordance, we also generated gene concordance factors (gCF) and site concordance factors (sCF) with IQ-TREE using the preferred maximum-likelihood topology as a reference. Gene concordance factors quantify the proportion of decisive gene trees supporting a given branch, whereas site concordance factors quantify the proportion of decisive alignment sites supporting that branch. Quartet support statistics (q1, q2, q3) together with branch lengths in coalescent units were extracted from ASTRAL analyses to evaluate the strength and distribution of conflict across the phylogeny.

As an alternative site-based multispecies coalescent approach, phylogenies were also inferred using CASTER-SITE V1.23.2.6 (Zhang, et al. 2025), which reconstructs species trees directly from site-quartet frequencies and therefore does not require prior gene-tree estimation. This approach provides an alternative assessment of species relationships, thereby avoiding potential biases arising from gene-tree estimation error. Analyses were conducted on amino acid and nucleotide concatenated datasets, including occupancy-filtered matrices (50%, 70%, and 75% thresholds), a dataset filtered according to variable-site occupancy, and a dataset excluding Neelipleona. CASTER-SITE branch lengths, quartet support values, and inferred topologies were compared with ASTRAL results to assess the consistency of species-tree estimation and phylogenetic conflict across alternative multispecies coalescent frameworks.

### Topology testing on concatenated datasets

Alternative hypotheses for the relationships among the four major collembolan orders (Entomobryomorpha, Poduromorpha, Symphypleona, and Neelipleona) were evaluated using likelihood-based topology tests implemented in IQ-TREE. All the fifteen fully resolved inter-order arrangements were represented as constrained backbone topologies. To obtain fully bifurcating candidate trees, each backbone constraint was resolved using the 70% occupancy amino-acid dataset (70pOccAA) under the LG+F+Γ model, which provided sufficient phylogenetic signal while minimizing missing data for reliable branch-length estimation. The resulting fifteen candidate trees were subsequently optimised and compared using site-wise likelihoods calculated from 70pOccAA.

Topology tests were conducted under LG+F+Γ and LG+C20-40-60+F+Γ to assess the sensitivity of hypothesis testing to model complexity and across-site compositional heterogeneity. Statistical support for competing hypotheses was evaluated using the approximately unbiased (AU), Kishino-Hasegawa (KH), Shimodaira-Hasegawa (SH), expected likelihood weight (ELW), and bootstrap proportion based on RELL (bp-RELL) tests with 10,000 RELL replicates. To further evaluate model robustness, all the topologies were re-analysed using the PMSF approximation (LG+C60+F+Γ+PMSF) on the top two topologies.

### Gene-wise topology testing

To investigate the distribution of phylogenetic signal among loci, topology tests were performed on individual genes meeting a minimum taxonomic representation of at least of three representatives from each collembolan order, and at least three of the four designated outgroup taxa. For each locus, the best-fitting substitution model identified during gene-tree inference was retained. For each locus the fifteen ordinal-level candidate hypotheses were first resolved into fully bifurcating topologies and subsequently pruned to retain only taxa present in the focal locus. Candidate topologies were pruned independently for each locus so that only taxa present in the alignment contributed to the likelihood calculations. Unlike previous studies, which evaluated only a limited number of alternative hypotheses, our framework exhaustively tested all fifteen fully resolved ordinal arrangements permitted for the four collembolan orders. Site-wise likelihoods were then calculated under the locus-specific substitution model and alternative topologies were compared using AU tests with 10,000 RELL replicates. Topologies with p-AU < 0.05 were considered significantly rejected, whereas the remaining hypotheses with p-AU ≥ 0.05 were considered not statistically rejected for that locus.

### Phylogenetic signal partitioning and progressive topology analyses

To quantify the ability of individual loci to discriminate among competing backbone hypotheses, loci were ranked according to the difference in log-likelihood between the best- and second-best-supported inter-order hypothesis (ΔlogL_best-second_). Alternative ranking schemes incorporating both ΔlogL_best-second_ and bp-RELL support were also explored. These rankings were used to assess whether support for alternative ordinal relationships was concentrated in a small subset of highly informative loci or distributed broadly across the dataset.

To examine how species-tree inference changed as progressively less informative loci were incorporated, loci were added cumulatively according to their ΔlogL_best-second_ ranking. Starting with the ten highest-ranked loci, additional loci were incorporated sequentially, and species trees were inferred using ASTRAL after each addition. Shifts in the recovered ordinal backbone were used to identify transition points between competing phylogenetic hypotheses. These transition points were then used to partition loci into discrete signal classes defined by changes in the inferred species tree. Loci belonging to each partition were concatenated and analysed independently using likelihood-based topology tests under increasingly complex substitution models (LG+F+Γ, LG+C20+F+Γ, LG+C40+F+Γ, and LG+C60+F+Γ). This approach allowed us to determine which subsets of loci preferentially supported alternative inter-ordinal relationships and whether the signal remained robust to increasingly realistic models of sequence evolution.

To quantify the concentration of topological support across alternative hypotheses, we computed the normalized Shannon entropy of bp-RELL weights for each dataset–model combination. Entropy was calculated as −Σ(p·log p)/log(N), where p represents the bp-RELL weight of each topology and N the number of topologies with non-zero weight, yielding values bounded between 0 (all support concentrated on a single topology) and 1 (support uniformly distributed across all alternatives).

### Molecular clock analyses

Divergence time estimation in Collembola remains particularly sensitive to fossil placement because the oldest known fossils predate the diversification of most extant lineages and often lack diagnostic characters that allow confident assignment to modern families or orders (Yu, et al. 2022). Consequently, different studies have adopted substantially different calibration strategies, resulting in markedly different evolutionary timescales. Leo et al. (2019) and Yu et al. (2024) calibrated several Palaeozoic fossils within extant collembolan lineages, including nodes associated with Entomobryomorpha and Isotomidae. However, the phylogenetic affinities of many of these fossils remain uncertain, and/or assigned before a comprehensive phylogenomic framework was available. To minimise the impact of uncertain fossil placement, we adopted a conservative calibration strategy.

Divergence times were estimated using MCMCtree V4.10 implemented in PAML (Yang 2007) under the approximate-likelihood framework (dos Reis, et al. 2015). Divergence-time analyses were performed using the preferred species-tree topology and the 70% occupancy amino-acid alignment filtered to retain variable sites (70pOccAAvsite). An independent-rates relaxed clock model was used, allowing evolutionary rates to vary among branches. Overall, fossil calibrations were based primarily on the calibration scheme of (Yu, et al. 2024), with some modifications to accommodate uncertainties surrounding the placement of early collembolan fossils.

Rather than assigning Paleozoinc fossils directly to extant collembolan lineages, we adopted a conservative calibration strategy that minimises uncertainty accociated to with fossil placement. Specifically, we constrained the root of Hexapoda and the crown of Collembola using broad age intervals that can be supported by the fossil records. In particular, *R. praecursor* (*incertae sedis*),, the oldest known collembolan fossil, was treated as a minimum age constraint for crown Collembola, rather than assigned to any specific extant order or family, while the oldest known pancrustacean fossil defined the maximum age (Uhen, et al. 2023). This strategy reduces the risk of over-constraining divergence times estimates through uncertain taxonomic assignments and allows molecular data to contribute more strongly to age estimation. Furthermore, our analyses incorporate substantially broader taxon sampling than previous studies and explicitly evaluates calibration interactions through prior-only analyses, providing an independent assessment of the robustness of the inferred timescale. Therefore, the root calibration (Hexapoda) was implemented as uniform prior, B(4.69,5.41), corresponding to 469-541 Ma. Crown Collembola was constrained with a uniform prior B(4.04,5.41) (404-541 Ma), using *R. preacursor* as minimum age. The remaining five internal calibrations were applied within Collembola following (Yu, et al. 2024). Because no Paleozoic fossil can be confidently assigned to any extant collembolan order, this calibration was applied to the crown group rather than to internal nodes. The remaining five internal calibrations followed Yu et al. (2024). These fossils were implemented as soft minimum constraints using gamma-distributed priors of the form G(2,0.35,*t*min), where *t*min corresponds to the fossil minimum age. This parameterisation concentrates prior density near the fossil age while allowing older divergence times when supported by molecular data. The birth-death process prior was specified as BDparas = 1 1 0. Priors on the overall substitution rate and rate variation among branches were specified using rgene_gamma = 2 20 and sigma2_gamma = 1 10, respectively. Five independent MCMC chains were run with burnin = 30,000, sampfreq = 500, and nsample = 20,000, corresponding to approximately 10 million iterations per chain.

Convergence was assessed by comparing the five independent runs in Tracer V1.7.1 (Rambaut, et al. 2018). Effective sample sizes (ESS) were verified to exceed 200 for all parameters. Posterior age estimates are reported as median ages with associated 95% highest posterior density (HPD) intervals. To evaluate the effective marginal priors induced by interacting calibrations, we repeated each MCMCtree analysis using usedata = 0, thereby sampling node ages from the prior without using sequence data. Prior and posterior marginal age distributions were compared for calibrated nodes and major collembolan divergences.

To evaluate the sensitivity of divergence-time estimates to uncertainty in inter-ordinal relationships, the complete dating analysis was repeated using the second-best supported backbone topology (T4; Leo et al 2019), corresponding to the second-best supported hypothesis recovered by our phylogenomic analyses. Unlike the preferred topology, T4 did not recover Poduromorpha as the earliest-divergence collembolan lineage. All calibration priors, clock settings, substitution-rate priors, and MCMC parameters were kept identical between analyses. Resulting divergent-time estimates and 95% HPD intervals were then compared between the two topologies.

## Results

### Taxon sampling, data filtering, and phylogenomic dataset construction

The final dataset comprised 145 collembolan taxa representing all four extant orders and 19 families, together with four outgroup species (Table S1). Orthology inference recovered 1,127 single-copy orthologous groups, representing the largest phylogenomic dataset assembled for Collembola to date.

We developed and evaluated an HMM-based filtering pipeline to mitigate the effect of hidden paralogy and spurious domain-level matches caused by extracting data from fragmented genome assemblies. Before the HMM-based filtering, orthologous groups showed a broad distribution of gene-species associations, consistent with substantial gene-to-species and species-to-gene noise (Fig. 2b, c). Following HMM-based filtering, this distribution collapsed into a markedly tighter distribution, indicating a substantial reduction in conflicting signal and improved orthology assignment, suggesting that many initial matches may represent paralogous copies or fragmented domain hits. Despite this pruning, broad taxonomic coverage was retained, and optimization converged on substantially higher compromise scores, consistent with improved agreement among loci and reduced phylogenetic conflict.

The impact of filtering was evident at both the locus and taxon levels. Across loci, the median proportion of species removed decreased from 31.3% to 17.3% (Wilcoxon signed-rank test, p = 2.2 × 10^−16^). The effect was particularly pronounced for loci identified as duplicated (putative paralogues), and subsequently split (partitioned), for which median removal decreased from 36.3% to 5.3% (p = 8.36 × 10^−14^). These loci accounted for a substantial fraction of the conflicting signal present in the initial dataset, indicating that hidden paralogy is a significant source of error in BUSCO-derived orthologue sets. A similar pattern was observed across taxa. The median proportion of genes removed per species decreased from 37.5% to 24.6% (Wilcoxon rank-sum test, p = 2.2 × 10^−16^), accompanied by a marked reduction in variance (Fig. 1b, c). This result suggests that a subset of lower-quality assemblies disproportionately contributed to paralogous signal and that HMM-based filtering effectively reduced this assembly-specific bias.

Taken together, these results demonstrate that hidden paralogues remain pervasive in BUSCO-derived datasets but can be substantially mitigated through HMM-guided filtering and locus partitioning. By improving both locus-level orthology and taxon-level consistency, this approach enables the inclusion of fragmented genomic resources while maintaining a robust phylogenetic signal.

### A robust phylogenomic backbone for Collembola

The final phylogenomic dataset comprised 145 collembolan taxa representing 19 families, together with four outgroup species, and included 1,127 single-copy orthologous loci. Gene occupancy was moderate overall, with species recovering an average of 663 loci (median = 735). Approximately 76% of loci were present in at least half of the taxa, whereas only 29% and 11% of loci were retained at the 70% (70pOccAA) and 75% occupancy (75pOccAA) thresholds, respectively. Individual loci averaged 822 nucleotides in length, with a median exceeding 1,000 nucleotides. Upon concatenation, the final amino-acid supermatrix (FullAAsmatrix) comprised 439,631 aligned positions, including 278,772 parsimony-informative sites.

Maximum-likelihood analyses of FullAAsmatrix recovered a highly resolved phylogeny with maximal or near-maximal support across most nodes (Fig. 3; Table S2). Nearly all families were recovered as monophyletic, except for Paronellidae, Neanuridae, and Hypogastruridae. All four extant collembolan orders were strongly supported as monophyletic. Support declined at the deepest backbone nodes, but all inter-order relationships remained supported by bootstrap values exceeding 75%, with the backbone topology placing Poduromorpha as sister to all remaining Collembola, followed by a divergence separating Entomobryomorpha from a clade comprising Symphypleona and Neelipleona.

**Figure 3.**
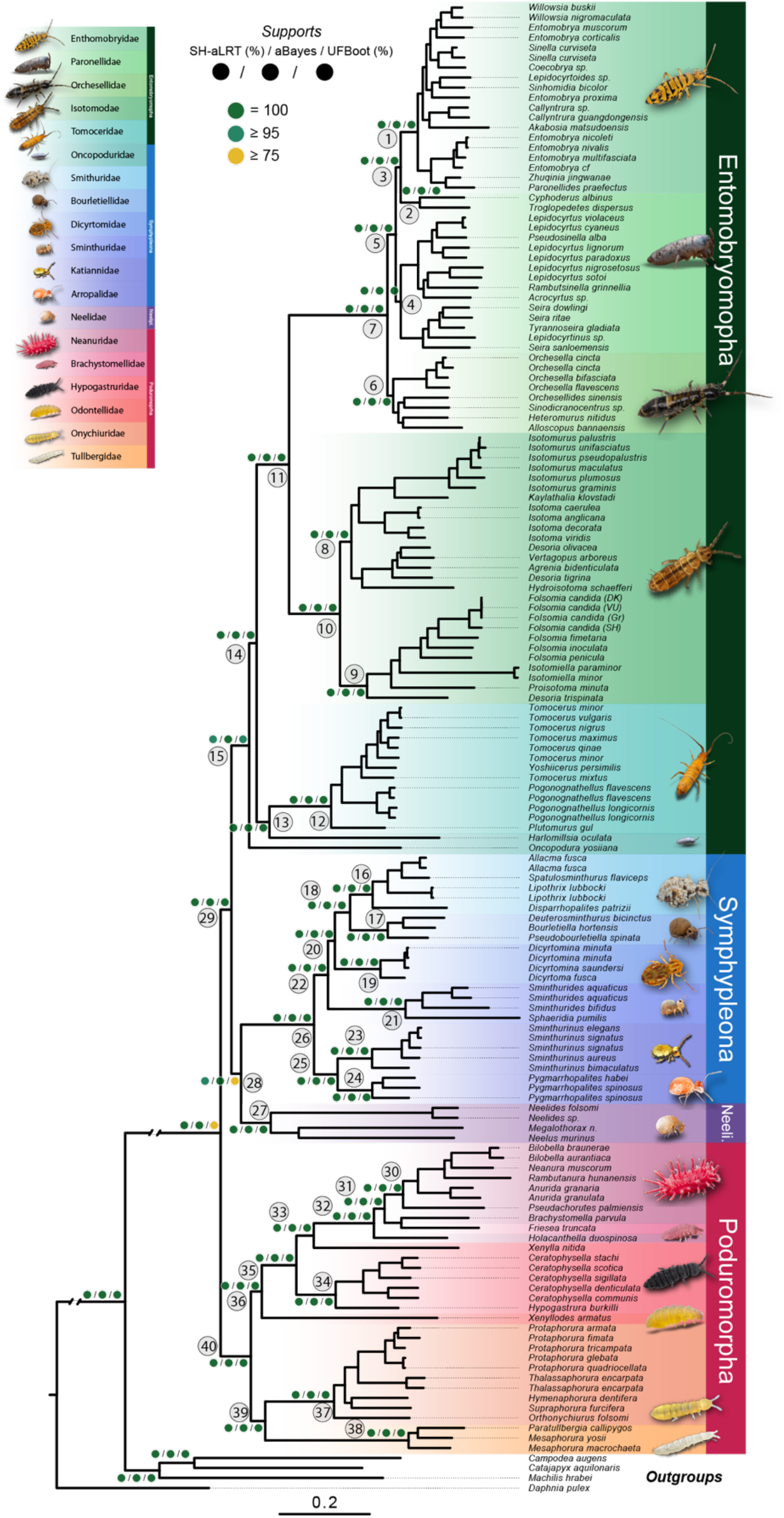
Maximum-likelihood phylogeny of Collembola inferred from FullAAsmatrix under the LG+F+Γ partitioned model comprising 149 taxa. Node support values are shown as SH-aLRT / aBayes / ultrafast bootstrap (UFBoot), with coloured symbols indicating support thresholds (green = 100%, dark green ≥95%, yellow ≥75%). Major collembolan orders are highlighted by coloured background bands: Entomobryomorpha (green), Symphypleona (blue), Neelipleona (purple), and Poduromorpha (red). Family-level and higher-level clades discussed in the text are numbered and correspond to the nodes evaluated across alternative phylogenetic analyses. Scale bar indicates amino-acid substitutions per site.

**Figure 4.**
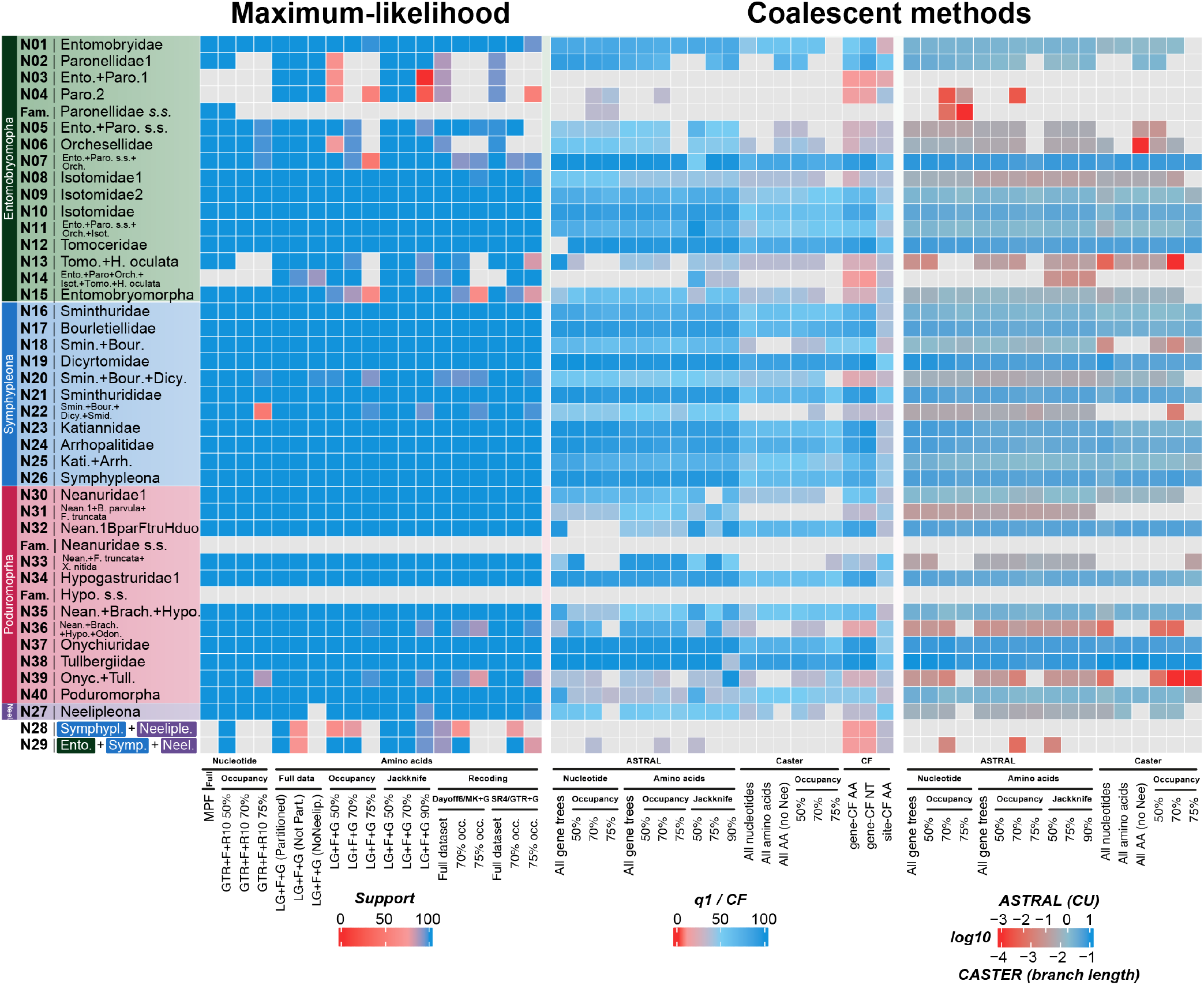
Concordance and robustness of phylogenetic relationships across alternative analytical frameworks. Heatmaps summarising support, concordance, and branch-length estimates for predefined family-, superfamily-, and order level clades across concatenation-, coalescent-, and site-based phylogenetic analyses. Rows correspond to the numbered clades shown in Figure 3 (full clade definitions and data are provided in Table S2-4), while columns represent alternative datasets and inference methods. Grey cells indicate that a clade was not recovered or that the corresponding statistic was unavailable for a given analysis. The left panel shows nodal support across maximum-likelihood analyses.

The full nucleotide supermatrix (FullNTsmatrix) recovered a largely congruent phylogeny, differing primarily in recovering Paronellidae as monophyletic, and in the arrangement of the backbone topology. In this analysis, Neelipleona were recorded as sister to all remaining collembolans, followed by Poduromorpha, sister to a clade comprising Symphypleona and Entomobryomorpha (Fig. S1). Despite this difference, support values for the deepest nodes remained moderate to high (SH-aLRT / UFBoot > 78%), indicating that alternative backbone arrangements are supported by the data rather than arising solely from poorly resolved nodes.

Across all sensitivity analyses, the amino-acid ML topology was recovered by the gene-jackknife replicates generated from 50%, 70%, and 90% random subsamples of loci and the majority of occupancy-filtered datasets. Alternative backbone arrangements were recovered only in 75pOccAA and in FullNTsmatrix (Fig. 3). To assess the potential effects of compositional heterogeneity and saturation, amino-acid alignments were recoded using the Dayhoff-6 and SR4 schemes. The majority of these recoded analyses recovered the same backbone topology as the full amino-acid analysis (full dataset Dayhoff-6, 70% Dayhoff-6 and 70% SR4 datasets). The remaining recoded analyses recovered either an alternative topology, which still retained Poduromorpha as the earliest-diverging lineage (75% SR4), or the same backbone topology inferred from the nucleotide datasets (full dataset SR4 and 75% Dayhoff-6; Fig. 3; Table S2). Collectively, these analyses indicate that two Poduromorpha-first topologies are consistently recovered across amino-acid datasets, jackknife replicates, recoding analyses, and taxon-exclusion analyses.

Collectively, these analyses narrow the uncertainty to the deepest relationships among the four extant collembolan orders, with amino-acid datasets consistently supporting two closely related Poduromorpha-first topologies and nucleotide datasets favouring a Neelipleona-first arrangement.

### Coalescent analyses reveal extensive gene-tree conflict

To evaluate whether the concatenation topology was robust to gene-tree discordance, we reconstructed species trees using ASTRAL and CASTER-site and quantified branch-specific concordance using gene concordance factors (gCF), site concordance factors (sCF), and ASTRAL quartet support. Although all methods recovered the four collembolan orders as monophyletic and largely agreed on family-level relationships, they disagreed on the relationships among the orders, revealing extensive gene-tree conflict concentrated at the deepest backbone nodes.

Full-dataset ASTRAL analyses recovered different backbone topologies depending on data type. The nucleotide gene-tree analysis placed Entomobryomorpha as the earliest-diverging order, followed by a clade comprising Neelipleona sister to Symphypleona + Poduromorpha. While the amino-acid gene-tree analysis recovered two sister clades, one comprising Neelipleona + Poduromorpha and another with Symphypleona + Entomobryomorpha (Fig. S1; Table S3). CASTER-site recovered yet other arrangements depending on data type. Using FullAAsmatrix, Symphypleona was placed as the earliest-diverging order, followed by Poduromorpha, with Entomobryomorpha and Neelipleona as sister groups, while FullNTsmatrix recovered the same backbone arrangement as the full nucleotide ML analysis (Fig. S1).

Analyses based on occupancy-filtered and jackknife datasets were broadly consistent with these patterns, with one notable exception, in three datasets (70pOccNT, 70pOccAA, and JK70pAA) ASTRAL recovered Poduromorpha as the earliest-diverging lineage, consistent with FullAAsmatrix ML topology, suggesting that in both data types, occupancy filtering partially recovers the ML signal under the coalescent framework. Notably, the backbone nodes supporting all these alternative arrangements were generally characterized by low quartet support (q1 ~ 35%) and short Coalescent Units (0-0.05 CUs). The only notable exception was the Neelipleona + Poduromorpha clade recovered from the full AA gene-tree dataset in ASTRAL, which exhibited a relatively high quartet support (q1 = 79%) despite low coalescent units (CU = 0.05).

To investigate the nature of these conflicts at backbone nodes, we examined the relationship between the dominant quartet frequency (q1) and CU across all nodes and ASTRAL analyses (Fig. S2). Under multi-species coalescence, these two measures are expected be positively correlated. Very low CUs should be associated with q1 values approaching one third, consistent with hard polytomies or rapid radiation that generated extensive incomplete lineage sorting (ILS), whereas high CU should be associated with high q1 values, reflecting stronger concordance among gene-trees and reduced ILS. Conversely, departure from this expected relationship, particularly nodes exhibiting relatively high CU but low q1 values, suggest that low concordance cannot be explained solely by insufficient time for lineage sorting, and instead may reflect genuine conflict among loci supporting reticulate evolution and alternative evolutionary histories.

Most order- and family-level nodes conformed to the expected pattern. Symphypleona consistently occupied the upper end of the distribution, with the highest q1 and CU values (q1s ~ 90%; log_10_CUs > 0), indicating a strongly supported and well-resolved phylogenetic signal. Neelipleona and Entomobryomorpha occupied intermediate positions (50% < q1s < 70%; log_10_CUs < 0), consistent with moderate resolution. Poduromorpha, on the other hand, showed a markedly anomalous pattern: relatively high CUs (log_10_CUs ~ 0) combined with very low q1 values (ASTRAL q1s < 50%; sCF = 31%), a combination inconsistent with either a hard polytomy or a well-resolved split, indicative strong conflicting signal among loci supporting alternative placements. This pattern was consistent across almost all ASTRAL analyses, except for FullNTsmatrix and JK50pAA, which showed intermediate q1 values for Poduromorpha (70 < q1 < 90), suggesting residual coherent signal in those subsets.

Collectively, these analyses failed to identify a single backbone topology consistently supported across coalescent frameworks. Instead, alternative inter-order relationships were associated with low quartet support and short internal branches, indicating that the deepest divergences in Collembola are characterized by pervasive gene-tree discordance.

### Topology testing identifies two statistically supported hypotheses for early collembolan diversification

To explicitly evaluate these topological conflicts, we tested all fifteen rooted topologies describing the possible relationships among the four collembolan orders. Using 70pOccAA and the site-heterogeneous LG+C60+F+Γ model, nine of the fifteen hypotheses (60%) were rejected by the approximately unbiased (AU) test (p-AU < 0.05), leaving six topologies not rejected by the AU test (Fig. 5a). Among these, the FullAAsmatrix ML topology (T11) consistently achieved the highest likelihood and received the largest bootstrap proportion based on RELL resampling (bp-RELL = 0.63). In terms of ΔlogL, the second-best topology was T3 (bp-RELL = 0.10), followed by T4 (bp-RELL = 0.13), and T2 (bp-RELL = 0.04). Notably, T3 and T2 both recovered Poduromorpha as the earliest-diverging collembolan lineage, whereas T4 corresponded to the topology previously proposed by Leo et al. (2019).

**Figure 5.**
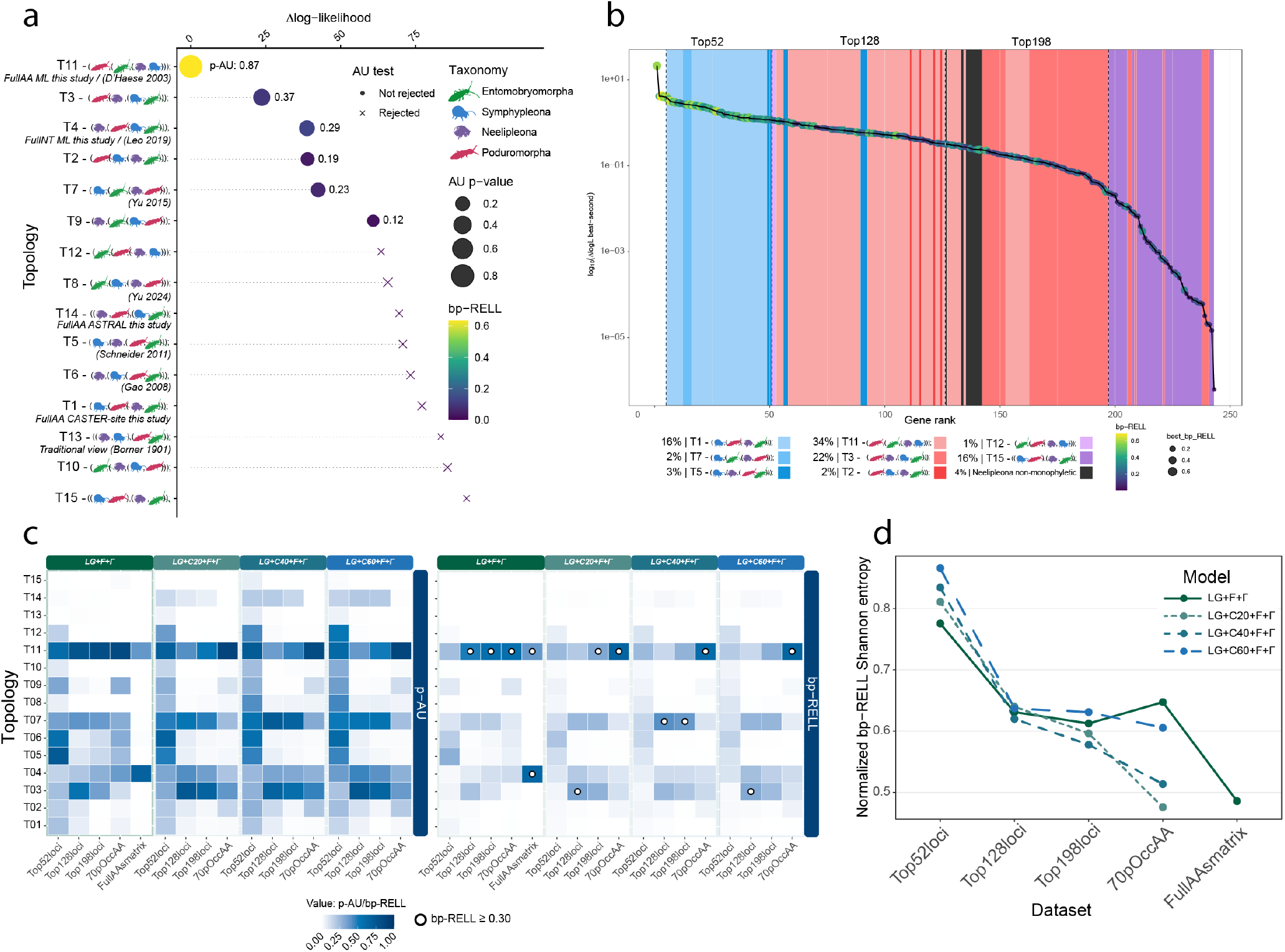
Gene-wise support for alternative backbone hypotheses and the effect of locus selection on ordinal-level relationships in Collembola. **a**) Comparison of 15 alternative backbone topologies (T1–T15) representing competing hypotheses for the relationships among the four collembolan orders. Topologies include the preferred concatenated topology recovered here (T11), the corresponding coalescent-based alternatives, and previously proposed arrangements from the literature. For each hypothesis, support is summarised by the difference in log-likelihood relative to the best topology (ΔlogL), approximately unbiased (AU) test results, AU p-values, and bp-RELL support. Schematic cartoons indicate the ordinal arrangement implied by each topology. **b**) Distribution of phylogenetic signal across the 243 loci used for topology testing, ranked by the difference in log-likelihood between the best- and second-best-supported topology for each gene. Vertical dashed lines indicate the cut-offs used to define the Top52, Top128, and Top198 subsets of the putative most topology-informative loci. Insets summarise the proportion of loci favouring the major competing backbone hypotheses among the ranked loci. **c**) Support for the 15 alternative topologies across the FullAAsmatrix and reduced locus subsets (Top52, Top128, Top198), and the 70pOccAA under increasingly complex substitution models. Heatmaps show AU p-values (left) and bp-RELL support (right) for each topology. **d**) Normalized entropy of bp-RELL support across the 15 candidate topologies for each dataset and model combination. Lower entropy indicates that support is concentrated on fewer competing hypotheses, whereas higher entropy indicates a more diffuse distribution of support among alternative backbones. Together, these analyses show that a relatively small subset of loci contains most of the ordinal-level phylogenetic signal, but that this signal remains distributed across multiple competing backbone hypotheses rather than converging on a single fully stable solution.

When the analysis was repeated using FullAAsmatrix under LG+F+Γ, support became stabilised on only two topologies. All alternative hypotheses were rejected, leaving T11 and T4 as the only arrangements not excluded by the AU test. Although T4 received a slightly higher bp-RELL value (0.60 versus 0.40 for T11), the two topologies differed by only 42 log-likelihood units despite being evaluated on a supermatrix containing more than 439,000 aligned amino-acid positions.

Collectively, these analyses reduced the original set of fifteen possible rooted topologies to only a small subset of supported hypotheses, many placing Poduromorpha as one of the earliest-diverging collembolan lineages.

### Distribution of backbone-discriminating signal across loci

To investigate the distribution of phylogenetic signal across loci, we performed gene-wise topology tests on the 243 most complete USCO loci using all fifteen alternative backbone hypotheses (see Methods). Gene-wise topology tests revealed that most loci contained insufficient information to discriminate among alternative inter-order hypotheses (Table S5). More than 57% of loci failed to reject even a single topology, and over 95% were unable to reject more than five alternative arrangements (Fig. S3).

To quantify the contribution of individual loci to backbone resolution, genes were ranked according to the difference in log-likelihood between the best- and second-best-supported topologies (ΔlogL_best–second_; Fig. 5b). This ranking revealed a skewed distribution of phylogenetic informativeness, with a relatively small subset of loci with a larger likelihood difference and a long tail with very little discriminatory signal. Cumulative analyses showed that approximately half of this signal was contained within the highest-ranked loci, whereas many genes contributed little or no information for resolving relationships among the four collembolan orders (Fig. 5b).

To test whether the deepest phylogenetic signal was concentrated in a small subset of highly informative loci, we generated a series of cumulative ASTRAL analyses by progressively adding loci according to their ΔlogL ranking. Rather than converging progressively toward a single topology, distinct backbone regimes emerged along the signal gradient. The highest-ranked loci (Top52) predominantly recovered topology T1, a hypothesis rejected by the concatenated topology tests. Inclusion of intermediate-ranked loci shifted support toward T11 and T3, the topologies favoured by concatenation-based analyses and topology testing. However, when all 243 loci were included, ASTRAL converged on the full coalescent topology (T15), indicating that conflicting backbone signal is not randomly distributed among loci. Instead, loci with different levels of topology-discriminating power contribute distinct and sometimes contradictory quartet signals, allowing the cumulative coalescent estimate to shift as progressively less informative loci are incorporated.

To further evaluate the distribution of phylogenetic signal, we partitioned loci into three groups corresponding to the major topology transitions observed in the cumulative analyses (Top52, Top128, and Top198). Each partition was concatenated and subjected to topology testing under increasingly complex substitution models ranging from LG+F+Γ to LG+C60+F+Γ. These datasets were analysed alongside 70pOccAA and FullAAsmatrix to test if backbone-discriminating signal was concentrated within a small subset of loci or progressively accumulated distributed across increasingly larger dataset.

Contrary to expectations that the highest-ranked loci would provide the clearest resolution, the signal-enriched Top52 dataset showed limited ability to discriminate among competing hypotheses. Even under the most complex models (LG+C40+F+Γ and LG+C60+F+Γ), support remained distributed across multiple topologies and no single arrangement exceeded bp-RELL values of 0.3 (Fig. 5c). As additional loci were incorporated, however, topology discrimination improved substantially regardless of model complexity. Across datasets and models, four topologies (T11, T7, T4, and T3) consistently retained support, with T11 repeatedly achieving the highest bp-RELL values. Importantly, highest-ranked loci were not markedly better at resolving backbone relationships than the larger 70pOccAA or FullAAsmatrix. A notable exception was topology T7 (the Yu et al. 2016 and Cucini et al. 2021 topology), which received transient support under intermediate mixture models (LG+C40+F+Γ) in highest-ranked loci but lost support under LG+C60+F+Γ.

To quantify the overall concentration of support across competing topologies, we calculated the normalised Shannon entropy from the bp-RELL values, for each dataset and model combination (Fig. 5d). High entropy indicates that support is broadly distributed among multiple topologies, and therefore contains little discriminatory information. In contrast, low entropy indicates that support is concentrated on a smaller number of competing alternatives. Entropy declined consistently from the smallest signal-enriched dataset (Top52) to the larger datasets (70pOccAA and FullAAsmatrix), while model complexity had a comparatively little or no effect. Together, these analyses indicate that the ability to discriminate among backbone hypotheses emerges progressively as information is accumulated across many loci, rather than being driven by a small subset of highly informative genes and model selection.

### Temporal origin of higher taxonomic groups and extant collembolan diversity

Divergence-time analyses placed the origin of crown Collembola in the Early Devonian, with a median age estimate of approximately 407 Ma (Fig. 6). The major extant orders diversified subsequently during the Carboniferous, with crown Poduromorpha, Entomobryomorpha, and Neelipleona originating between approximately ~346 Ma and ~300 Ma, whereas crown Symphypleona originated considerably later, during the Late Triassic (~221 Ma; Table S7).

**Figure 6.**
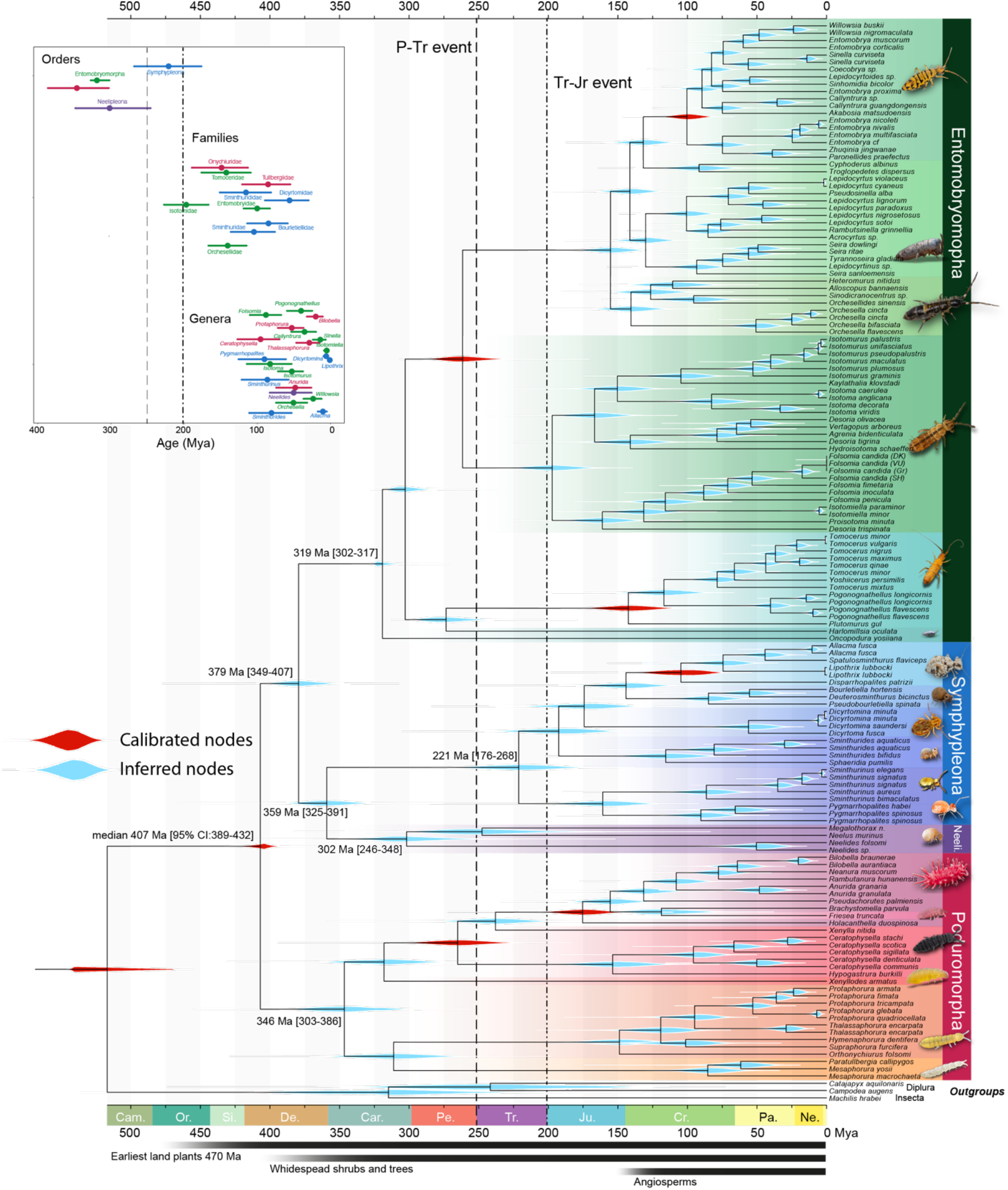
Time-calibrated phylogeny of Collembola. MCMCtree under the preferred maximum-likelihood topology. Red density plots indicate fossil calibration priors, whereas blue density plots represent posterior age estimates for inferred nodes. Horizontal bars correspond to 95% highest posterior density (HPD) intervals. Geological periods are shown along the time axis. The inset summarizes crown-age estimates for sampled orders, families, and genera, with points indicating posterior median ages and horizontal bars representing 95% HPD intervals.

To assess calibration behaviour, MCMCtree was run without sequence data to sample from the joint prior distribution. Posterior age estimates differed markedly from the marginal priors for most calibrated nodes, indicating that molecular data, rather than calibration priors, primarily informed divergence-time estimates (Fig. S4). Then, to estimate the sensitivity of divergence-time estimates to deep backbone uncertainty, the dating analysis was repeated using topology T4 (Leo, et al. 2019), the second-best topology recovered in topology testing. Despite the alternative backbone topology, divergence-time estimates remained remarkably stable (Fig. S5; Table S7). Median age estimates differed by less than 3.3 Myr for the Collembola crown, Entomobryomorpha, Symphypleona, and all examined family-level clades. Larger differences were restricted to the lineages directly affected by the backbone rearrangement, namely Neelipleona (59 Myr) and Poduromorpha (19 Myr).

The temporal distribution of crown ages revealed a clear hierarchical pattern across lower taxonomic ranks. Most extant families originated primarily during the Jurassic and Cretaceous (Fig. 6; S6). Amongst the sampled families, Isotomidae represented one of the oldest crown groups (~200 Ma), followed by Orchesellidae (~140 Ma), Sminthuridae (~105 Ma), Entomobryidae (~100 Ma), Bourletiellidae (~85 Ma), and Dicyrtomidae (~55 Ma). Most remaining families originated within the same broad interval, indicating that a substantial fraction of modern family-level diversity accumulated during the Mesozoic, broadly comparable to patterns reported for many other terrestrial hexapod groups.

Remarkably, many extant genera possess crown ages that overlap with those of numerous families, indicating an unusually deep temporal persistence of genus-level lineages. For example, the cosmopolitan genera *Folsomia, Isotoma, Sminthurinus, Pygmarrhopalites* or *Ceratophysella* are estimated to have originated more than 75 Ma ago, during the Late Cretaceous or earlier, comparable to or older than several currently recognized families (Fig. 6; S7). Suggesting that extant collembolan genera represent ancient evolutionary lineages that have accumulated diversity over tens of millions of years.

## Discussion

The phylogenomic analyses presented here recover a well-resolved and broadly stable phylogenetic framework for Collembola above the family level, yet inter-order relationships remain stubbornly challenging to resolve. Importantly, this uncertainty does not appear to arise from insufficient data or poor taxon sampling. Instead, multiple lines of evidence suggest that the early diversification of extant collembolan orders occurred over a very short evolutionary interval, which left weak and conflicting phylogenetic signals distributed across genomes. Understanding the origin of this conflict and its implications is a central challenge in elucidating the evolution of the earliest terrestrial hexapods (Du, et al. 2024, Du, et al. 2025, Machida, et al. 2025).

### Amino acid and nucleotide datasets recover different backbone topologies

One of the clearest outcomes of our analyses is the systematic discordance between amino acid and nucleotide datasets. Whereas most analyses of amino-acid datasets recovered a Poduromorpha-first topology (*e*.*g*.: T11), an arrangement not previously proposed by molecular studies, but consistent with the morphology-based hypothesis of (D’Haese 2003), nucleotide datasets consistently supported a Neelipleona-first arrangement (T4), in line with earlier molecular analyses (Leo, et al. 2019, Sun, et al. 2020a). This is not surprising *per se*, because deep time divergences are particularly susceptible to substitutional saturation, compositional heterogeneity, and lineage-specific biases in codon usage, which accumulate over hundreds of millions of years, disproportionally influencing nucleotide-based analyses. Translation into amino-acids reduces the impact of synonymous substitutions and partially mitigates these biases, often improving performance at ancient phylogenetic depths (Liu, et al. 2014). Consistent with this interpretation, Dayhoff-6 and SR4 recoding analyses largely recovered the same backbone relationships as the original amino-acid datasets despite dramatically reducing the amino-acid state space. Furthermore, support for T11 increased also with model complexity were adopted LG+F+Γ to LG+C60+F+Γ, suggesting that realistic modelling of among-site compositional heterogeneity favour the Poduromorpha-first hypothesis.

Nevertheless, recoding analyses did not eliminate all uncertainty, and some recoded datasets recovered topologies resembling those obtained from nucleotide analyses (SR4 GTR+Γ). These observations suggest that systematic error alone cannot explain the observed conflict. Instead, the disagreement between amino acid and nucleotide datasets likely reflects a combination of model inadequacy, saturation, and genuine biological discordance generated during the earliest phases of collembolan diversification.

### Concatenation and coalescent approaches disagree at backbone nodes

One of the major sources of discordance involves the contrast between concatenation-based and coalescent-based approaches. Concatenated amino-acid analyses consistently favoured T11, whereas ASTRAL and CASTER-site recovered multiple alternative backbone arrangements. Such disagreements are increasingly recognised in ancient radiations and are often associated with extensive incomplete lineage sorting (ILS) and reticulate evolution (Suh, et al. 2015).

Several observations suggest that this explanation is relevant here. Internal coalescent branch lengths at the backbone nodes are extremely short, typically ranging between 0 and 0.05 coalescent units. At the same time, gene-wise topology tests revealed that most loci contain little information capable of discriminating among alternative backbone hypotheses. More than half of all loci failed to reject a single topology, and over 95% were unable to reject more than five alternatives. Under these conditions, species-tree methods operate on collections of gene trees that are themselves only weakly informative regarding the deepest divergences.

The cumulative ASTRAL analyses provide additional insights. Rather than converging toward a single solution, species-tree estimates shifted repeatedly as progressively less informative loci were incorporated (Fig. 5b). Yet topology testing and entropy analyses revealed the opposite pattern: support became increasingly concentrated on a small number of hypotheses as additional loci were added. This result suggests that the deepest phylogenetic signal is not concentrated in a handful of highly informative genes; instead, support for particular backbone arrangements emerges gradually through the accumulation of many individually weak signals distributed across the genome.

An important observation is that topology testing did not distribute support evenly amongst alternative hypotheses, instead the highest-likelihood topologies consistently recovered Poduromorpha as the earliest-diverging collembolan lineage. Three of the four best-scoring hypotheses identified by AU testing shared this feature despite differing in the relationships among the remaining orders. Thus, much of the residual uncertainty concerns the branching order of Neelipleona, Symphypleona, and Entomobryomorpha rather than the position of Poduromorpha itself.

### The unusual phylogenetic signal of Poduromorpha crown

Within the four collembolan orders, Poduromorpha occupies a uniquely anomalous position. While Symphypleona, Neelipleona, and Entomobryomorpha generally display the expected positive relationship between quartet support and CUs, Poduromorpha consistently combines relatively low quartet support with unexceptionally high CUs. This pattern differs from that expected under a simple lack of phylogenetic information and instead suggests substantial conflict among loci.

One possibility is that the divergence leading to Poduromorpha occurred during a brief interval near the origin of crown Collembola, followed by a long period of lineage diversification and ILS, maybe with a reticulate evolution. Such scenario would generate a weak and heterogeneous signal at the ancestral split while allowing substantial CUs to accumulate subsequently.

Historically, various poduromorphan taxa, such as Tullbergidae have been associated with long-branch attraction artefacts, especially in association with Neelipleona (Sun, et al. 2020b). However, our results suggest this is not the case: excluding Neelipleona did not substantially alter backbone relationships, and recoding analyses continued to recover Poduromorpha-first topologies despite strong reductions in compositional heterogeneity. Taken together, these results suggest that the unusual behaviour of Poduromorpha reflects genuine properties of the underlying evolutionary history rather than simple analytical artefacts, corroborating previous morphological results (D’Haese 2003).

### A soft rather than hard polytomy

The combination of low quartet support and short coalescent branches indicates substantial conflict among alternative quartet resolutions, a pattern consistent with rapid early diversification in which successive lineage splits occurred over a short evolutionary interval, leaving limited opportunity for complete sorting of ancestral polymorphisms.

Overall, this evidence supports a scenario of rapid but resolved diversification rather than a true hard polytomy. Under a strict hard polytomy, alternative quartet resolutions should occur at approximately equal frequencies and all backbone hypotheses should remain statistically indistinguishable. Neither prediction is supported by our analyses. Although several backbone nodes show low quartet support, AU tests reject most of the fifteen possible ordinal arrangements, and support becomes progressively concentrated on a smaller subset of hypotheses as both locus number and model complexity increase.

The interpretation is reinforced by the distribution of bp-RELL across datasets. Shannon entropy constantly declines from the Top52 dataset to progressively larger matrices, largely irrespectively of model complexity (Fig. 5d), indicating that support becomes increasingly concentrated on fewer competing hypotheses. If the earliest collembolan divergences represented a true hard polytomy, increasing dataset size would not be expected to produce this pattern. Instead, the results are more consistent with a soft polytomy, in which successive divergences occurred over a geologically brief interval followed by a definite branching sequence, recoverable in principle. Uncertainty surrounding early collembolan diversification is therefore confined to a limited set of alternative backbone arrangements rather than spread across the full range of topologies proposed in the literature.

### Parallels with other ancient terrestrial radiations

The pattern documented here closely resembles that observed in several other ancient rapid radiations. Similar combinations of extensive gene-tree conflict, short internodes, and disagreement between concatenation and coalescent methods have been reported for early placental mammals, Neoaves (Suh, et al. 2015), major land-plant lineages (Wickett, et al. 2014), molluscan classes (Chen, et al. 2025), and deep arthropod divergences (Su, et al. 2024). In many of these systems, increasing genomic sampling has not eliminated conflict but instead revealed that alternative histories are genuinely represented among different regions of the genome.

Collembola occupy a particularly important position within this broader context. As one of the earliest confirmed terrestrial hexapod groups, their diversification was intimately associated with the establishment of Palaeozoic soil ecosystems. Our divergence-time estimates place the initial radiation of crown Collembola within the broader interval of terrestrial ecosystem expansion during the Devonian and Carboniferous, between ~390-430 Ma, with a median of ~400 Ma, just before the earliest forests originated. The extremely short coalescent internodes recovered among the four extant orders are therefore consistent with a rapid burst of diversification associated with the early assembly of terrestrial food webs.

Although uncertainty remains regarding the precise branching order of the four collembolan orders, multiple independent analyses converge on a limited set of plausible hypotheses, with Poduromorpha-first topologies receiving the strongest overall support. The persistence of alternative signals across analytical frameworks likely reflects the rapidity of the original radiation itself, whose genomic signature remains detectable more than 350 million years after the diversification of the earliest extant springtail lineages.

## Conclusions

Using the largest phylogenomic dataset assembled for Collembola to date, we provide a substantially revised framework for understanding the early evolution of springtails. Multiple independent analyses revealed that this uncertainty does not arise from insufficient data. Instead, concatenation, coalescence, concordance, topology-testing, and signal-partitioning approaches consistently indicate that the earliest diversification of crown Collembola was characterised by extensive conflict among loci and sites. Although alternative backbone arrangements were not rejected, the majority of analyses converge towards a small subset of alternatives, which hints at Poduromorpha being the most likely earliest-diverging extant lineage, whilst the remaining uncertainty is concentrated in the relationships amongst Entomobryomorpha, Neelipleona, and Symphypleona.

The distribution of phylogenetic signal across loci further demonstrates that support for deep collembolan relationships is not confined to a small number of highly informative genes. Rather, backbone resolution emerges through the accumulation of weak but consistent signal distributed across hundreds of loci. The persistence of conflict across analytical frameworks, combined with extremely short coalescent branch lengths and low quartet support values, is consistent with a reticulated evolution typical of rapidly evolving radiations accompanied by extensive incomplete lineage sorting.

Finally, our divergence-time analyses place the origin of crown Collembola within the early Devonian. This is broadly coincident with the expansion of early terrestrial ecosystems, and suggests that the diversification of the major extant orders occurred during the Carboniferous period. These estimates support a scenario in which springtails diversified during the early origins of terrestrial soil communities, making them amongst the oldest terrestrial arthropod radiations known today.

Collembola joins a growing list of ancient radiations in which genome-scale datasets reveal that deep phylogenetic conflict can persist despite extensive taxon and locus sampling. Rather than representing a limitation of phylogenomics, this conflict likely preserves a biological signature of rapid diversification more than 350 million years ago. Resolving the remaining uncertainty will require not only larger datasets, but also continued advances in evolutionary models capable of extracting signal from some of the oldest and most complex radiations in the history of terrestrial life.

## Acknowledgments

The authors are grateful to the computational facilities of the Advanced Computing Research Centre, University of Bristol. FC thanks Dr. Callum Wright, for his valuable HPC support, and Dr. Mattia Giacomelli for early discussion that helped motivate this work. We also thank the anonymous reviewers for their constructive comments, which improved the manuscript.

## Data Availability

Trees, datasets, and custom scripts can be found on Zenodo at ….

